# A new method to isolate algal species from mix algal culture

**DOI:** 10.1101/233981

**Authors:** Ehsan Ali, Saima S. Mirza

## Abstract

To meet the issues of energy and environment, algae cultivation for biofuel and CO_2_ sequestration is getting popular at the global level. Specific algal strains have been identified for production of biofuel, biomolecules and biomass. To start algae cultivation at lab or industrial scale, it is requirement to have isolated and identified algal culture for targeted products. Water sample for algae from aquatic system is usually consist of mix culture of algae and need to be processed for targeted isolated algal strains using reported techniques like streaking, spraying, serial dilution, and single-cell isolations. But none of these techniques is considered as efficient or popular except streaking on agar plate which involves a set of microbial techniques and may take months to make isolation properly. Here, a new method is proposed to make alginic acid solution using aquatic sample followed by pouring it in calcium chloride solution drop by drop which makes the beads with single or more algal species trapped in each bead. The trapped algal species in the beads are grown in 96 wells plate having single bead in each well with standard medium leading to microscopic verification of the isolated algal species to process further. A mix culture from a lake was subjected to isolation using proposed method and excellent results were obtained in one week duration.

## 1. Introduction

Advancement in microalgae derived biofuel and other targeted bio-products at lab scale or industrial level is contributing well towards management of fuel and environmental crisis for better human life on this planet. The scientists focusing on algal biofuel technologies may need some innovation in the prevailing algal strains isolation techniques which were reported first time about 60 years before. The techniques like streaking, spraying, serial dilution, and single-cell isolations are considered as time taking and uncertain towards quick isolation of potential microalgae candidate for lipid or other targeted products production. Doan et al, 2011 has used automated flow cytometric cell-sorting technique for isolation of algae which is complicated and expensive in comparison to the proposed technique for algal strain isolation [1]. The reported isolation techniques in practice are laborious and need extra ordinary skills to follow the procedure. Currently, four major techniques including streaking, spraying, serial dilution, and single-cell isolations are in practice at lab or industrial level. Most of these techniques have been reported about 50 to 70 years back. Allen and Stanier (1968) have reported a method to isolate blue green algae from water and soil samples based on temperature [2]. Gerloff et al. (1950) have reported that the streaking for isolation of algae on agar plates is unsatisfactory methods of separating blue-green algae from mix culture [3]. Currently, these techniques have been utilized by scientists working on algae or algal products with some modifications. Lee et al (2014) have reported isolation using streaking technique [4]. The whole process of isolation may take weeks to months for proper isolated algal strains Thus in the present study, an attempt was made to develop a novel, effortless, more applied and cost effective method to isolate the algal strains from aquatic samples using alginic acid beads to trap the scattered algal species in different beads and then regrow it in separate wells of 96 well ELISA microplate. Alginic acid, a naturally occurring hydrophilic colloidal polysaccharide is frequently used in drug delivery system [5]. Few reports are also available using alginic acid or alginate for heavy metal removal from wastewater [6]. This novel method represents a valuable addition for the development of an indigenous source of biolipid for biodiesel feedstock.

## 2. Methodology

### 2.1 Sample Collection

Water samples having prominent algal growth were collected from different natural water reservoirs of Kalar Kahar that is located in Chakwal District of Punjab, Pakistan. Sampling was done from multiple sites of large bodies. All samples were collected in 50 ml falcon tubes, transferred to laboratory and stored at 4°C in refrigerator for further use.

### 2.2 Sample Enrichment

For isolation of algae from mixed culture, collected samples were enriched by using multiple recipes for each sample. Recipes of nutrient media included BG 11 [7-8] and bold basal medium [9]. Enriched algal cultures in these media were mixed thoroughly and confirmed presence of multiple species under light microscope (Euromax).

### 2.3 Isolation and maintenance of unialgal culture

After microscopic study of the mix culture to ensure the presence of different algal strains, it was subjected to isolation using alginic acid and calcium chloride. The methodology involves making 10 ml 10% solution of sodium alginate (alginic acid) in distilled water and mixing it with 10ml of water sample of algae having mix culture giving 5% final concentration of sodium alginate. A 100ml solution of calcium chloride (0.2 M) was prepared using distilled water. A 5ml syringe was used to fill the algae plus alginic acid solution and then the same syringe was fixed with 17 G needle to pour the algal mixture prepared with alginic acid drop wise into 0.2 M calcium chloride solution, the beads were formed and kept in 0.2 M calcium chloride solution for 5-6 hours to strengthen calcium bonding for longer shelf life. There was a formation of 1-1.5 mm round beads possessing algal cells on random basis (Fig. 2). The beads were treated using three different procedures at ambient conditions as below:

The beads were transferred to 96 wells plate having 200 μl BG11 medium in each well and one bead per well. The beads were smashed partially using autoclaved tooth picks and then plate was kept in light and allowed to continue growth in the wells. After 3-5 days, the growth was observed in different wells. The slides were prepared using grown culture from wells and checked under microscope (Euromax) using high power (40 and 100 X).

## 3. Results and discussion

The mixed algal culture collected from lake was subjected to light microscopy visualization to ensure the presence of single or multiple algal strains to be processed further for isolation. A mixture of different algae was seen under high power as shown in Fig. 1. The mix algal culture or aquatic sample was contained of some thread like and round shaped algal species. The beads prepared using alginic acid and mix culture were supposed to be containing different algal cells on random basis (Fig. 2).

**Fig. 1.**
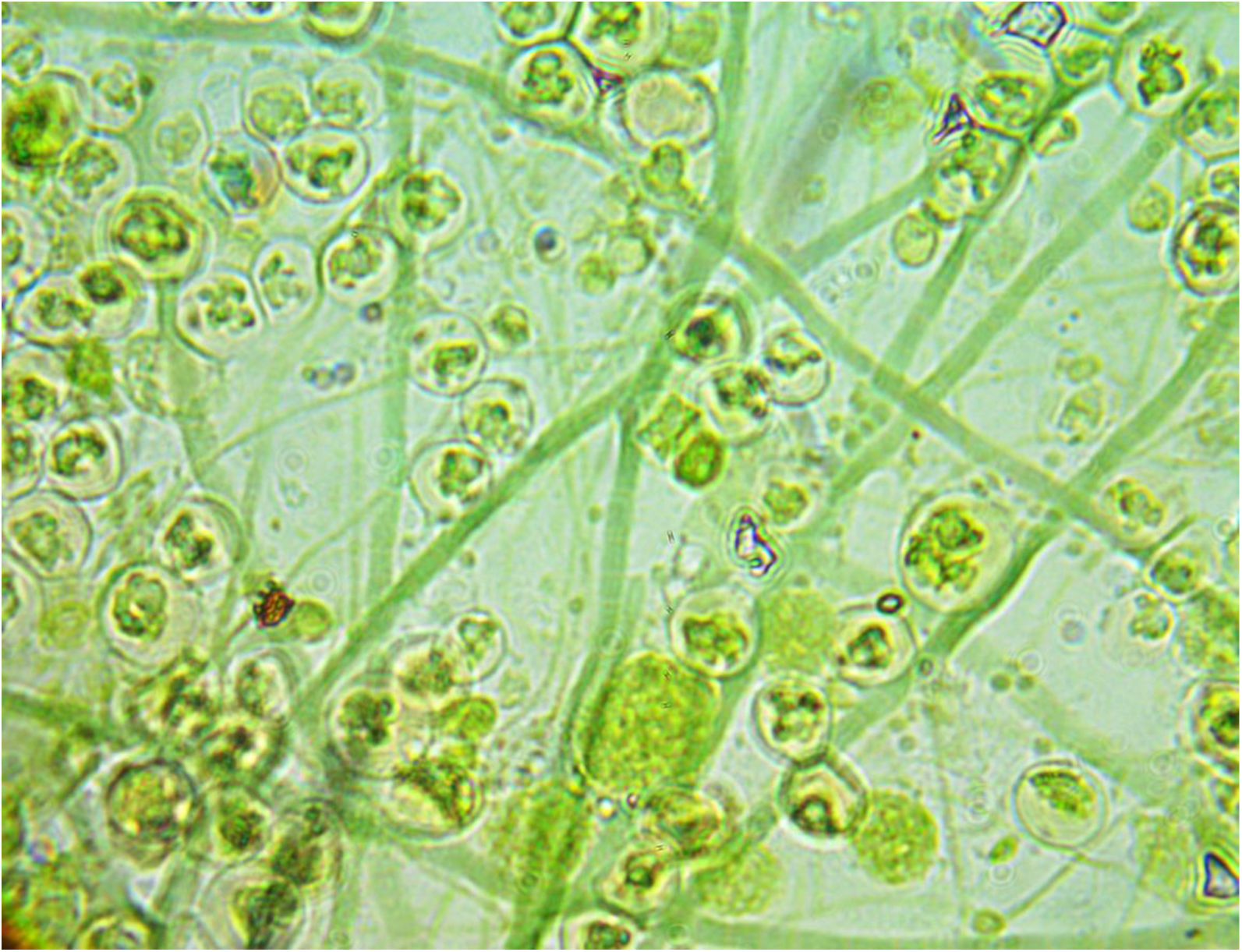
The mix culture of algae.

**Fig. 2.**
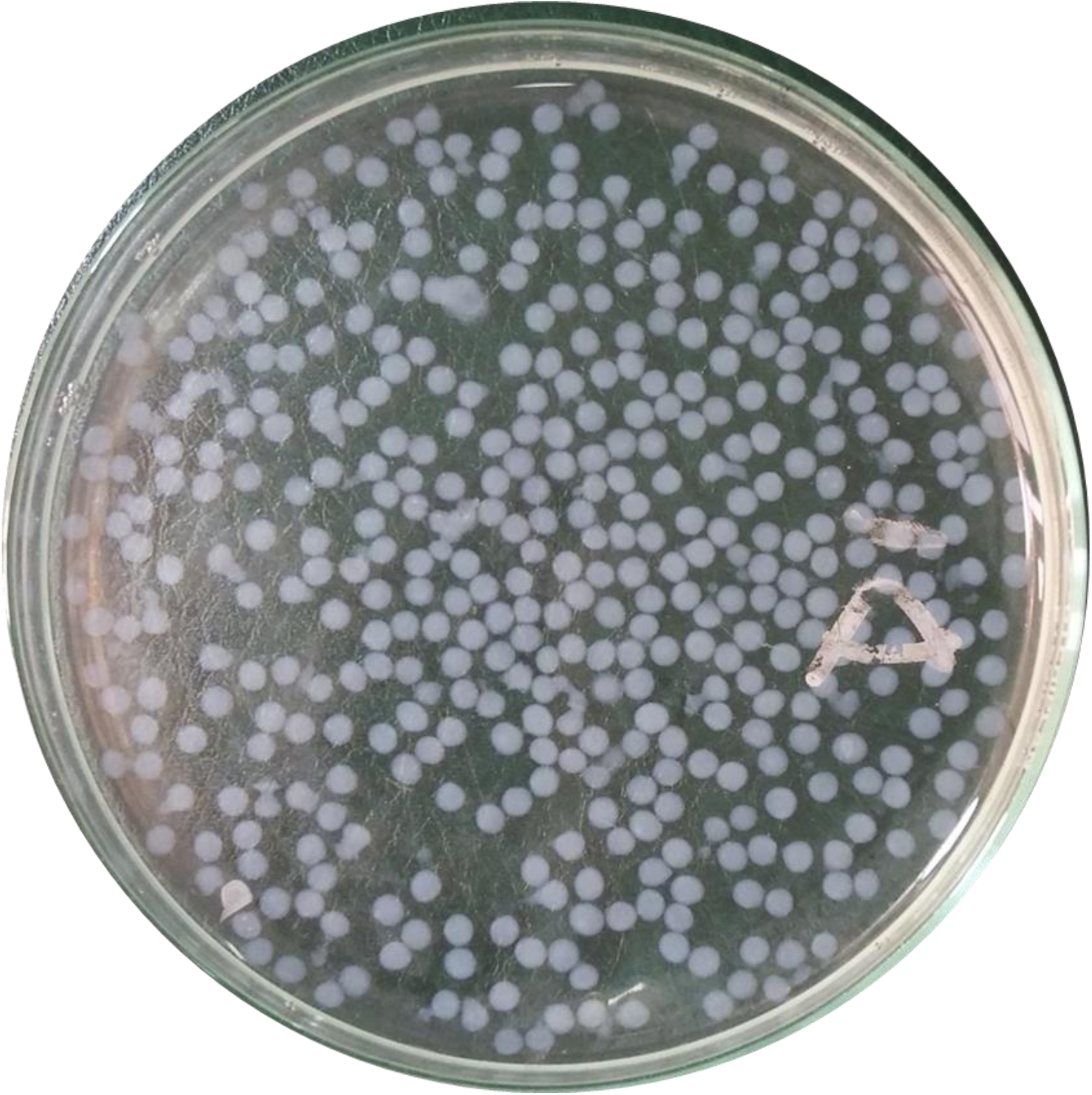
Alginic acid beads with embedded algal cells.

### 3.1 Isolation of different algal species by alginic acid beads

The beads prepared using mix culture and grown in isolated wells had shown growth in 3-5 days in few wells but after 7-8 days the growth was observed in almost all wells. There was no visible inhibitory effect on the growth of algae due to alginic acid or calcium chloride solution. The isolation mechanism in the present study involves trapping of algal cells into beads of alginic acid at random basis. After bead formation, the trapped algal species have exposure to surrounding and can use light and carbon dioxide for growth in the presence of nutrients as growth medium. Cancela et al. (2016) has reported coagulation of algal culture using calcium chloride, sodium alginate and tannins of *Eucalyptus globulus* bark but not for isolation of different strains from mix culture [10]. In addition to this, the calcium is a component of algal medium and may not have any inhibiting effect at appropriate concentration.

Fig. 3A shows the pouring or culturing of possible algal species embedded in the beads in different wells of 92 well plate. Each well was containing 200 μl of appropriate medium (BG-11) and exposed to light for photosynthetic activity of embedded algal cell/s in the beads. The growth started in 3-5 days in few beads or wells and after 7-8 days the growth was dense and appeared in most of the wells as shown in Fig. 3B. The exposure to light and access to nutrients to the embedded algal cells may vary depending upon the thickness of the bead and take varying time duration to respond for growth. Alginic acid or alginate has been widely used in drug delivery system and specifically to release the drug slowly for specialized metabolic activities in the body [11]. The characteristic of alginic acid described by the author as a slow drug releasing agent could be a factor to allow the growth of different algal species at different time duration. The variable time duration towards growth response by the embedded algal cells is in agreement with the report published by Kurczewska et al. (2014) [11]. The grown culture from 20 wells was used to make 20 slides and visualized under high power microscope. The seven slides out of 20 have shown isolated cultures. The three different algal species were found in this algal mixture and will be subjected to identification in further studies.

**Fig. 3.**
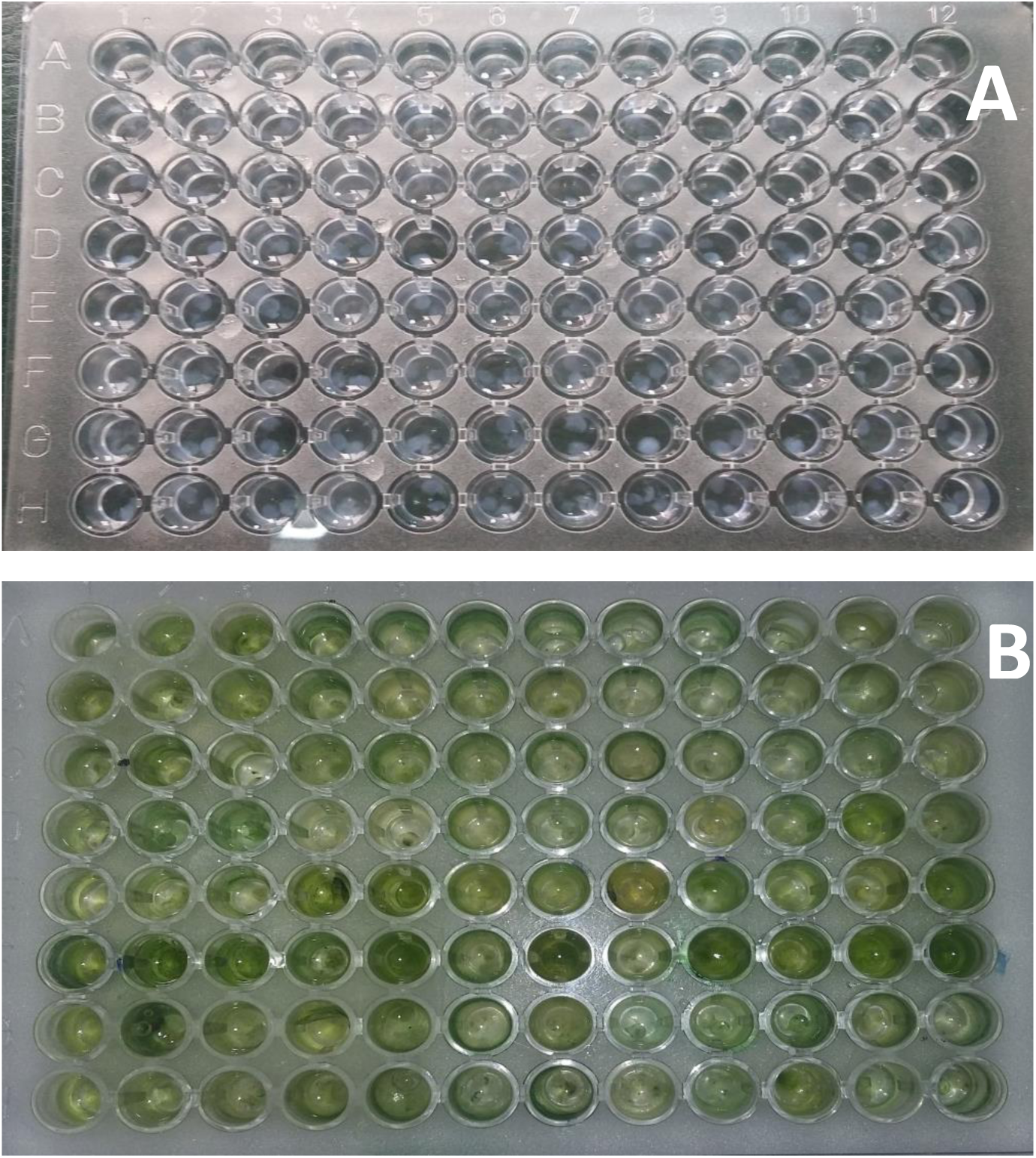
Alginic acid beads in 96 well plate with embedded algal species before growth (A) and after growth (B).

### 3.2 Morphological comparison of algal species isolated by serial dilution and alginic acid beads

Fig. 4–8 shows the clear pictures of isolated algal strains by alginic acid beads, respectively designated as algal specie (i-v) from the mix culture and may be processed further to get axenic culture leading to lab scale or mass cultivation of algae. Morphological comparison under microscope of three algal species isolated by alginic acid beads with their respective specie available in literature revealed clear morphological similarity.

**Fig. 4.**
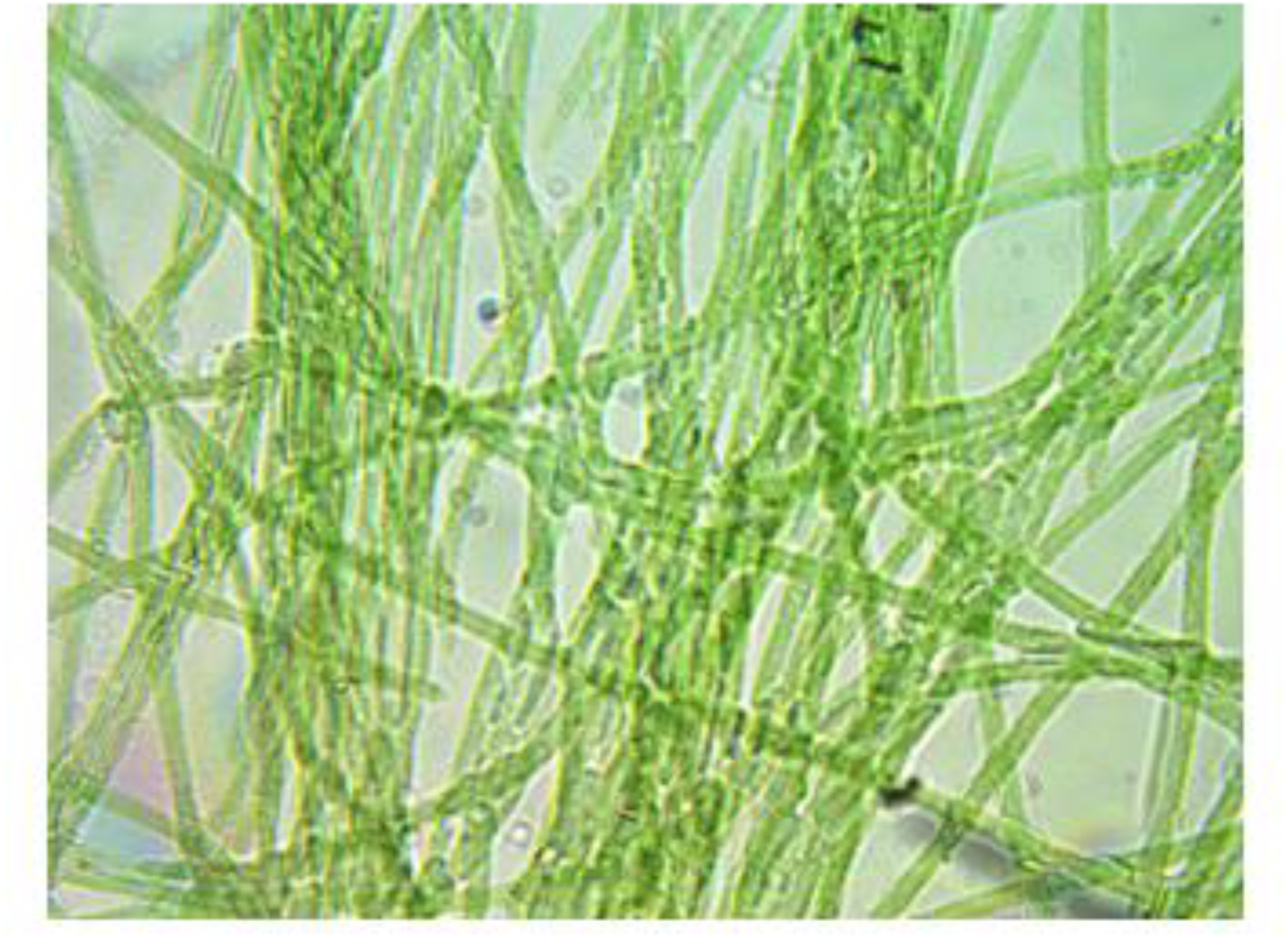
Isolated algal specie (i)

**Fig. 5.**
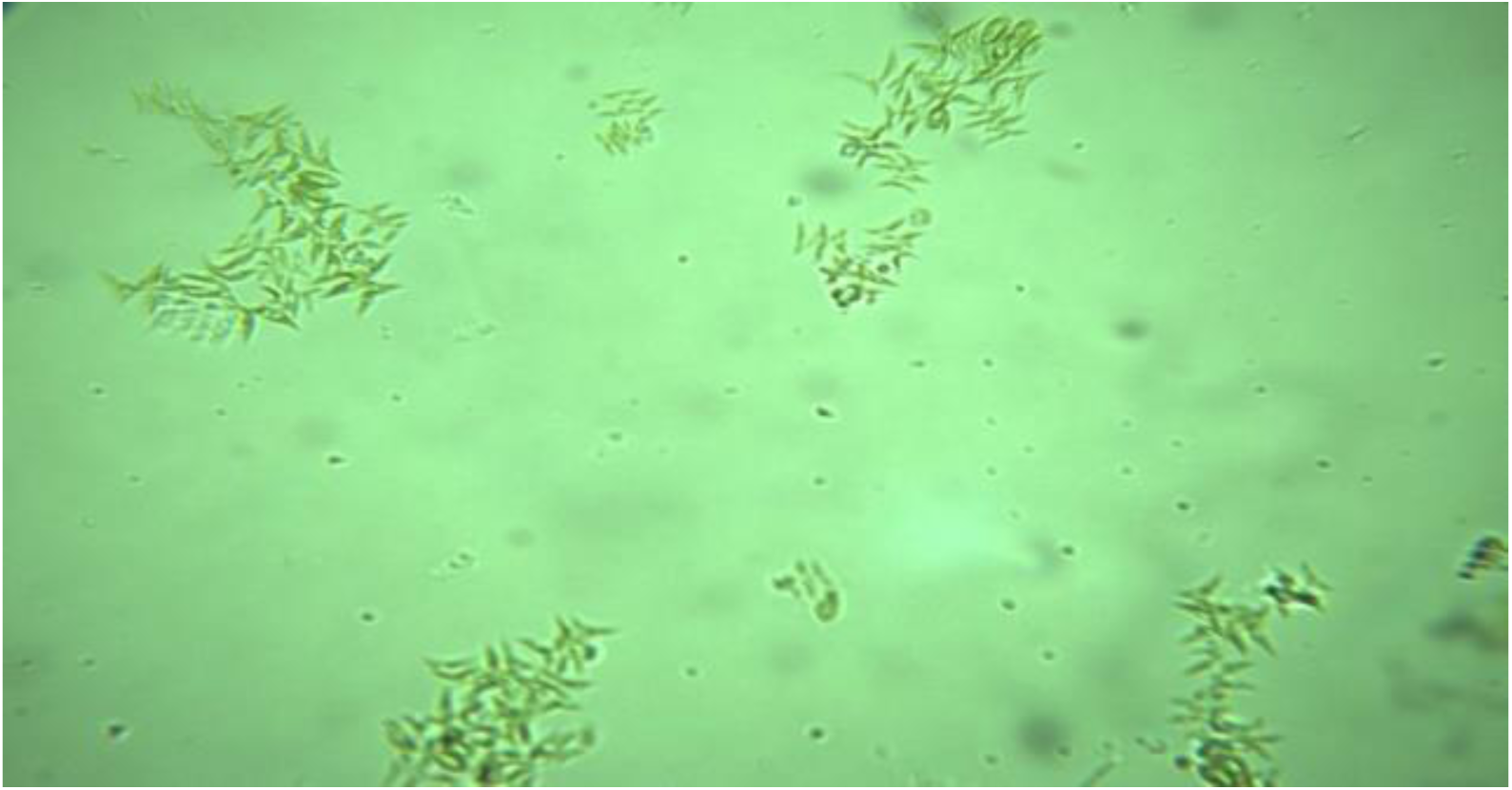
Isolated algal specie (ii)

**Fig. 6:**
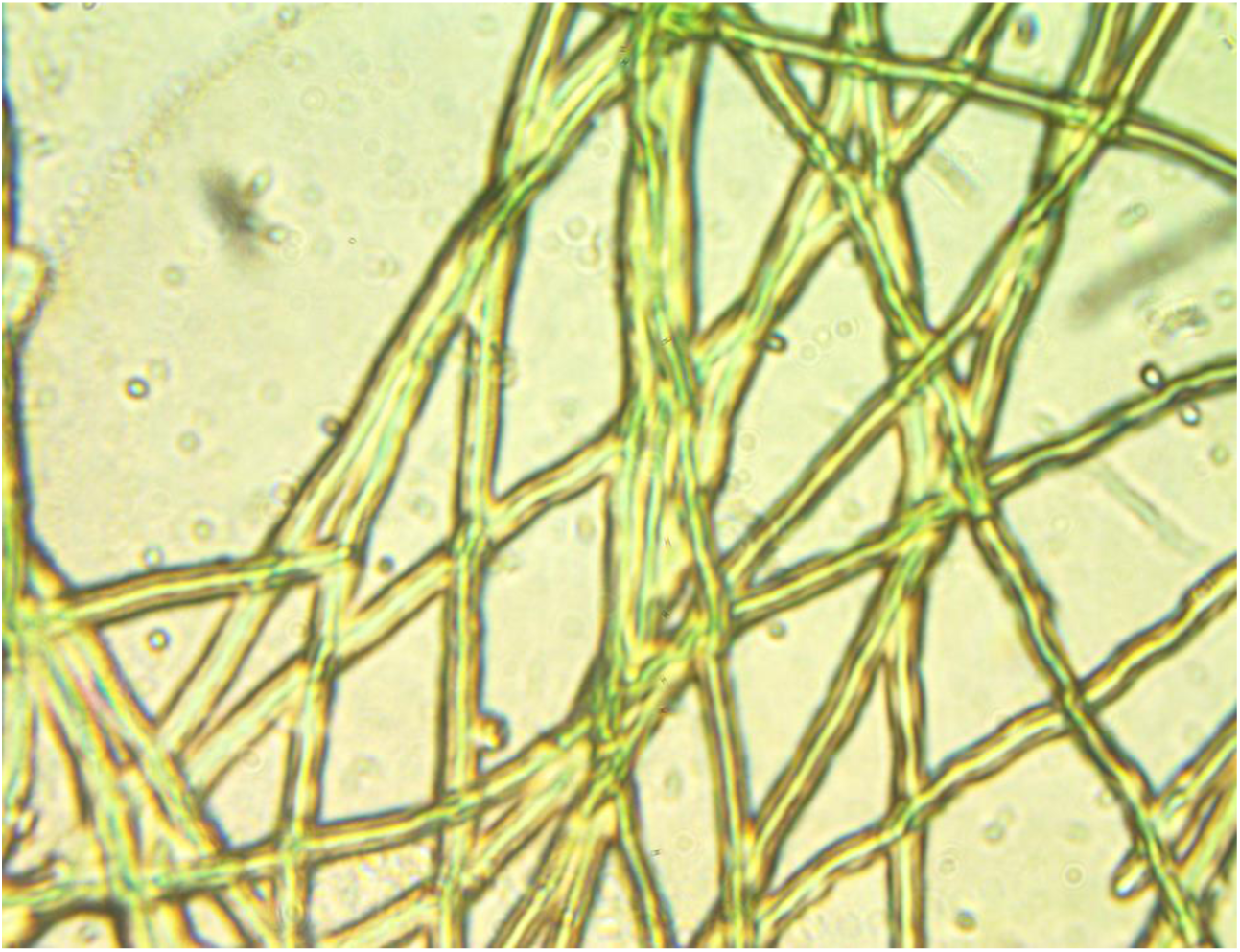
Isolated algal specie (iii)

**Fig. 7:**
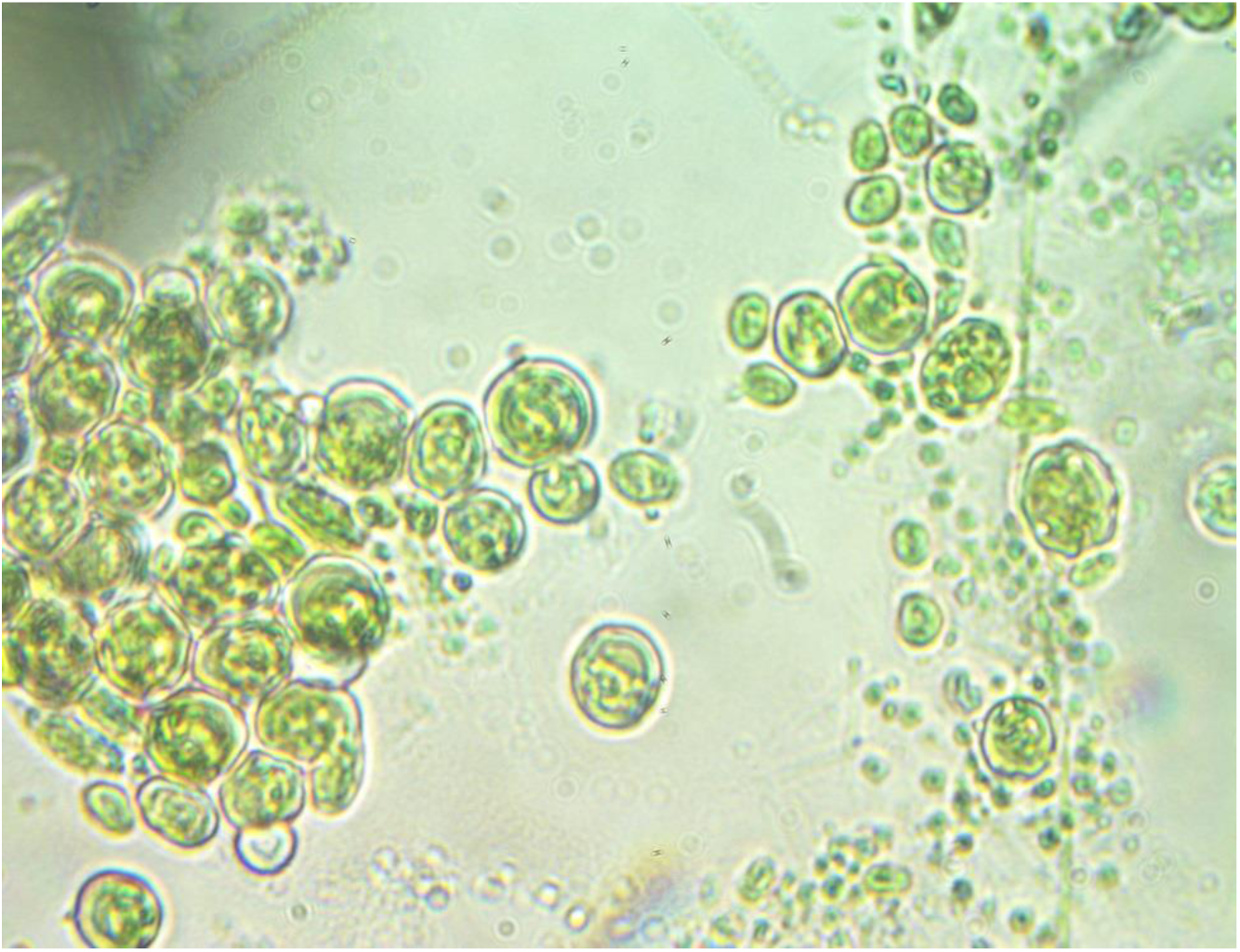
Isolated algal specie (iv)

**Fig. 8:**
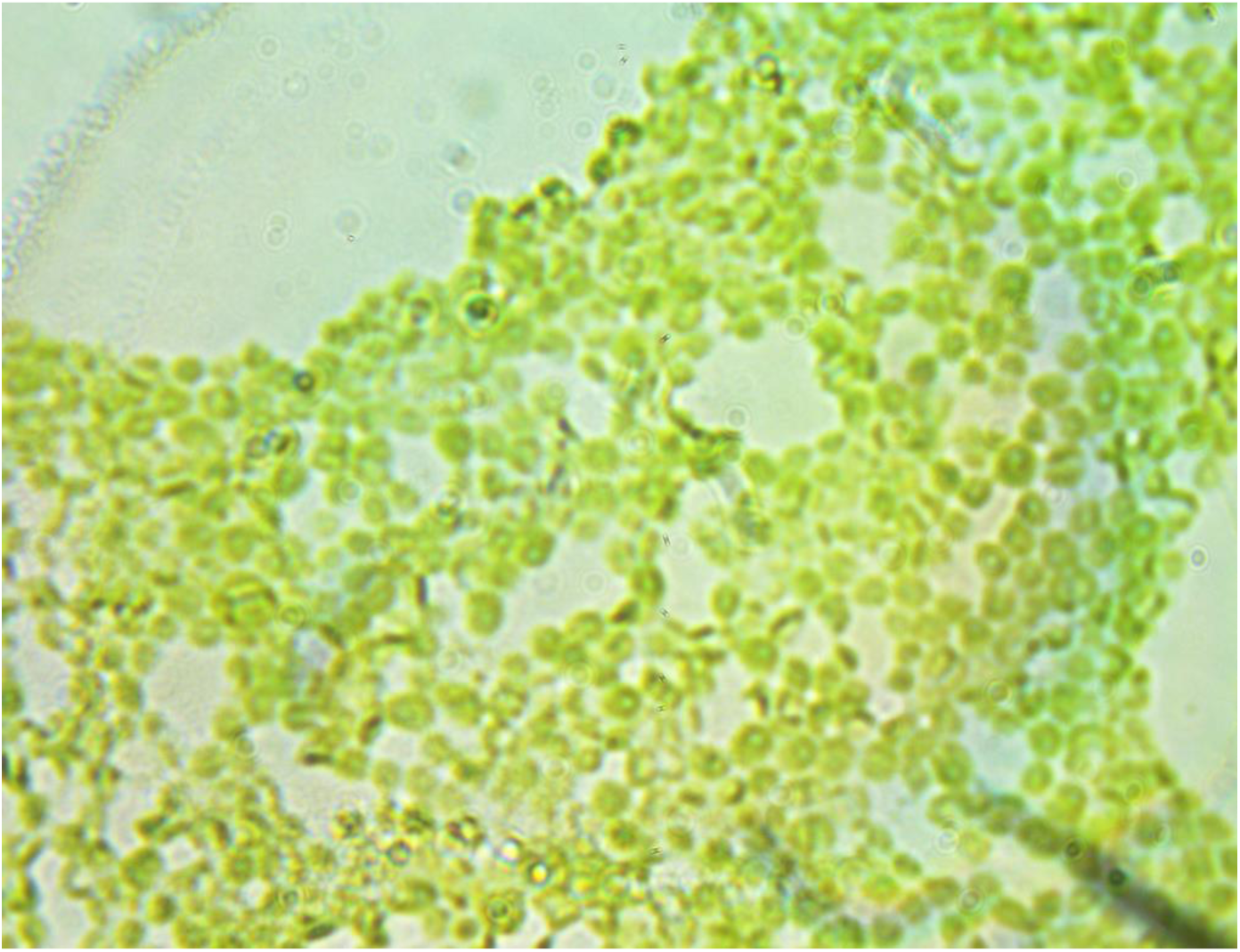

The technique presented here has several advantages to conventional techniques like streaking, serial dilution and micropipette washing technique. The streaking involves sterilization steps using autoclaving and spreading or streaking on petri dishes having agar medium then growth leading to selection of isolated algal colonies in separate tubes or flasks. The isolation of algal species using streaking may take months to get isolated algal strains and sometimes got contaminated by heterotrophic bacteria on agar medium with algal spreading and streaking. This contamination is probably due to organic contents constituted in agar medium [12]. Gerloff et al (1950) have reported that the isolation of blue green algae using streaking on agar plates is not satisfactory technique. But there was no work reported by any scientist later to improve the algal isolation techniques. Similarly, serial dilution also time taking and laborious technique. It may require subsequent dilution and may also be harmfully affected by contaminations [13]. Regarding micropipette washing technique, isolation and contamination free culture maintenance is not quite easy. During micro pipetting technique, it was experienced that identifying and picking axenic colonies or filaments is most critical in the whole process of algal isolation. Effective implementation of these techniques still entails considerable expertise and persistence. Henceforth, declining in the number of associated heterotrophic bacterial contamination is one of the key advantages of algal isolation technique presented here.

Nonetheless, other investigators have also reported different innovative methods for isolation of different algal species. For instance, Ferris and Hirsch et al. (1991) introduced a novel method for cyanobacteria isolation by employing nutrient rich glass fiber filters. Furthermore, the author employed an antibiotic for limitation of heterotrophic bacterial contamination. Doan et al., (2011) reported flow cytometric cell-sorting technique for screening of potential oil rich microalgae candidates would serve as biodiesel feedstock.

In a nutshell, all existing techniques for algal isolation lack simplicity and require energy input in terms of sterilization, subsequent dilutions, extensive centrifugation, instrumentation etc. The proposed technique is simple and can give accurate outcomes for isolation of algal strains from mix culture.

## 4. Conclusion

The proposed method for isolation of algal strains from mix culture or aquatic samples is an easy approach to make isolation of algae for different applications like mass cultivation of specific species for biofuel, biomolecules and biomass production. The use of alginic acid and calcium chloride is not presenting any inhibitory effect on the growth of embedded algal cells in standard algae medium. The mechanism involves the calcium bonding of alginic acid with embedded single or multiple algal cells forming into whitish beads making different algal cells separate from each other, and regrowth of these cells into separate containers or micropipette plate wells. This technique is more efficient and quick to get results with certainty as compared to conventional techniques reported about 70 years back.

## Acknowledgement

The authors are thankful to the Govt. of the Punjab, Pakistan for funding. The experimental assistance from students including Ms. Asma Andleeb, Ms. Sidra Akbar and Ms. Rubab is also acknowledged here

